# A hybrid *de novo* genome assembly of the honeybee, *Apis mellifera*, with chromosome-length scaffolds

**DOI:** 10.1101/361469

**Authors:** Andreas Wallberg, Ignas Bunikis, Olga Vinnere Pettersson, Mai-Britt Mosbech, Anna K. Childers, Jay D. Evans, Alexander S. Mikheyev, Hugh M. Robertson, Gene E. Robinson, Matthew T. Webster

## Abstract

**Background:** The ability to generate long sequencing reads and access long-range linkage information is revolutionizing the quality and completeness of genome assemblies. Here we use a hybrid approach that combines data from four genome sequencing and mapping technologies to generate a new genome assembly of the honeybee *Apis mellifera*. We first generated contigs based on PacBio sequencing libraries, which were then merged with linked-read 10x Chromium data followed by scaffolding using a BioNano optical genome map and a Hi-C chromatin interaction map, complemented by a genetic linkage map.

**Results:** Each of the assembly steps reduced the number of gaps and incorporated a substantial amount of additional sequence into scaffolds. The new assembly (Amel_HAv3) is significantly more contiguous and complete than the previous one (Amel_4.5), based mainly on Sanger sequencing reads. N50 of contigs is 120-fold higher (5.381 Mbp compared to 0.053 Mbp) and we anchor >98% of the sequence to chromosomes. All of the 16 chromosomes are represented as single scaffolds with an average of three sequence gaps per chromosome. The improvements are largely due to the inclusion of repetitive sequence that was unplaced in previous assemblies. In particular, our assembly is highly contiguous across centromeres and telomeres and includes hundreds of *AvaI* and *AluI* repeats associated with these features.

**Conclusions:** The improved assembly will be of utility for refining gene models, studying genome function, mapping functional genetic variation, identification of structural variants, and comparative genomics.

## Background

A complete and accurate genome assembly is a crucial starting point for studying the connection between genome function and organismal biology. High quality genome assemblies are needed for reliable analyses of comparative genomics, functional genomics, and population genomics [1]. High-throughput short-read sequencing technologies now allow the routine generation of massive amounts of sequence data for a fraction of previous costs [2]. Despite this, however, these data are not amenable to producing highly contiguous *de novo* assembly and tend to result in highly fragmented assemblies due to the difficulty in assembling regions of repetitive DNA sequence [3]. Many available genome assemblies, therefore, have low contiguity and are fragmented in repetitive regions [1]. Chromosomal structures of fundamental importance to genome function such as centromeres and telomeres are also rich in repetitive DNA and often missing from genome assemblies, which hinders studies of their role in cell division and genome stability. Repetitive sequences are also often involved in generating structural variants, which are important for generating phenotypic variation, and are implicated in processes such as speciation, adaptation and disease [4–7].

Several long-range sequencing and scaffolding technologies have been developed recently that can be used to produce *de novo* assemblies with hugely improved quality and contiguity [8]. The chief advantage of these technologies lies in their ability to span low-complexity repetitive regions. Here we utilize four of these methods: PacBio, 10x Chromium, BioNano and Hi-C. Pacific Biosciences (PacBio) SMRT sequencing uses single-molecule real-time sequencing to produce reads of tens of kilobases, enabling assembly of long contigs [9]. The linked-read 10x Genomics Chromium technology uses microfluidics to localize multiple short reads to the same molecule, facilitating scaffolding of short reads [10]. The BioNano optical mapping technology detects the occurrences of small DNA motifs on single molecules, which enables long-range scaffolding of assembled contigs [11–13]. The Hi-C method identifies chromosomal interactions using chromosome conformation capture that can be used to group and scaffold contigs using their physical proximity in the genome [14,15].

Each of these technologies suffers from weaknesses and no single technology alone is likely to generate an optimal assembly. For instance, assembly of long reads is still problematic in long highly-repetitive regions and it is challenging to generate sufficient depth across most eukaryotic genomes to produce chromosome-length contigs using long-read sequencing due to the long length of some repetitive regions and the sequencing cost [16]. Linked-read sequencing provides a significant improvement in contiguity over assemblies produced by short-read sequencing alone, but still suffers from the same drawbacks for assembling highly repetitive regions into complete contigs. Long-range scaffolding technologies such as BioNano are able to produce highly contiguous scaffolds, but it can be problematic to place short contigs on these scaffolds due to lack of homologous motifs [17]. Due to these various drawbacks, the current state-of-the-art for genome assembly is to use a hybrid approach combining multiple technologies [18–21]. Several genome assemblies produced in this fashion are of comparable or better quality than finished human and model organisms that have undergone large number of improvements with additional data [1,22–25].

The Western honeybee *Apis mellifera* is a species of huge importance to agriculture and ecology and a model for understanding the genetic basis of behavior and the evolution of sociality [26–29]. With the use of chromosome banding techniques, telomere- or centromere-labeling fluorescent probes, and genetic maps, the honeybee karyotype was well-established decades ago [30–33]. The honeybee genome is ~250Mbp and consists of one large metacentric chromosome with two long chromosome arms (chr. 1) and 15 smaller submetacentric/acrocentric chromosomes (chr. 2-16) [33], in which the centromere is located off-center and delineates a short and a long arm. The first published genome assembly (Amel_4.0), based on whole-genome shotgun sequencing with Sanger technology [33], suffered from poor coverage of low-GC regions and recovered unexpectedly few genes. An upgrade incorporating next-generation ABI SOLiD and Roche 454 sequencing of DNA and RNA (Amel_4.5), improved sequence and gene coverage [34], but the assembly was still fragmented (N50=0.046 Mbp) and large-scale features and repeats such as centromeres and telomeres were still largely missing or poorly assembled. An improved genome assembly is therefore of great utility for uncovering the function of genes and other chromosomal features.

Here we used four complementary technologies to generate a highly contiguous *de novo* assembly of the honeybee. We used closely related haploid drones in our analyses, which do not suffer from ambiguities in resolving heterozygous variants seen in diploid genomes. Our pipeline involved assembly of PacBio long read data into contigs, which were then merged and scaffolded with 10x Chromium linked-read data. Finally, we performed long-range scaffolding using BioNano optical mapping and Hi-C proximity ligation data. We describe extensive improvements in completeness and contiguity of this assembly compared to previous genome assemblies.

## Results

### Contig generation with PacBio and 10x Chromium

We generated data with PacBio, 10x Chromium, BioNano, and Hi-C. The PacBio and 10x Chromium sequences were first used to produce separate independent assemblies using FALCON and Supernova respectively (see Methods). The PacBio assembly had the highest contiguity of these single-technology assemblies, with 429 primary contigs of average size 0.520 Mbp and N50 of 3.09 Mbp (67 times longer than Amel_4.5; **Table 1**). We next scaffolded the PacBio assembly with 10x data using ARCS [35] and LINKS [36], and oriented contigs and scaffolds on a genetic map followed by additional gap filling with PBJelly [37]. The contiguity of this assembly version (Amel_HAv1) was significantly improved compared to both the individual 10x and PacBio assemblies (8.8-fold and 1.7-fold increase in N50, respectively). The longest Amel_HAv1 contig is 13.4 Mbp, 40 times longer than in the longest contig in Amel_4.5. N50 of the HAv1 is 5.167 Mbp, compared to 0.046 Mbp for Amel_4.5 (112-fold improvement; **Fig 1**; **Table 1**).

**Table 1.**
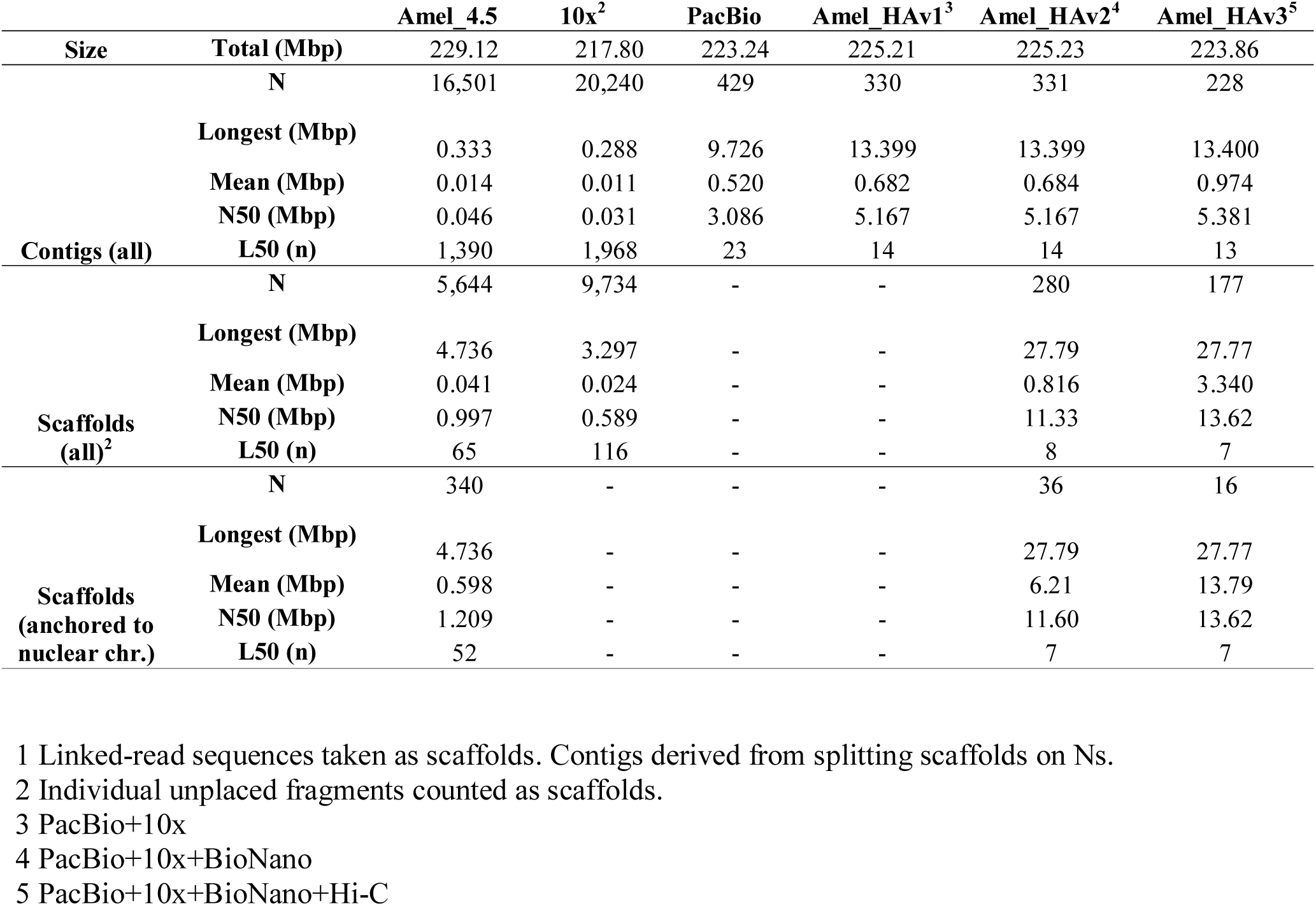
Overall assembly statistics

**Fig 1.**
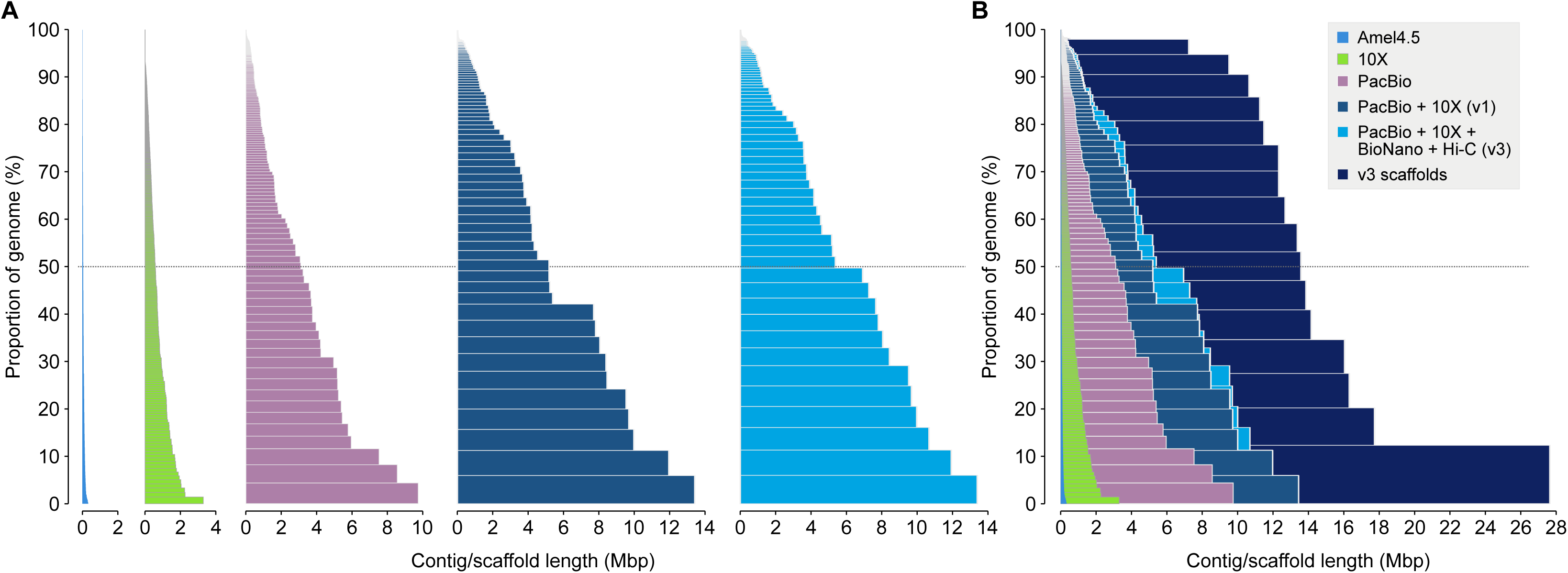
Comparison between assemblies. A) Stacked contigs from the official honeybee genome assembly Amel_4.5 [34] and the long-read sequencing technologies used in this project. Sequences are sorted by length (x-axis) and the cumulative proportion of each assembly that is covered by the contigs is displayed on the y-axis. Dashed line indicates contig with length equivalent to N50. From the left: Amel_4.5, 10x Chromium-only (assembled using Supernova), PacBio-only (assembled using FALCON), Amel_HAv1 (PacBio contigs + 10x scaffolding, see Methods) and Amel_HAv3 (Amel_HAv1 scaffolded using BioNano to produce AmelHA_v2, followed by Hi-C scaffolding). For 10x Chromium sequences, the full-length linked-read scaffolds are shown (i.e. including gaps). B) Stacks from A super-imposed over the Amel_HAv3 scaffolds (i.e. including gaps). These scaffolds are chromosome-length and contain 51 gaps.

### Scaffolding with BioNano and Hi-C

We performed scaffolding of the Amel_HAv1 contigs using BioNano data to produce version Amel_HAv2. This version contains 26 hybrid scaffolds with N50 of 11.3 Mbp and the longest scaffold of 27.8 Mbp. In total 96 out of 171 BioNano genomic maps and 77 out of 328 Amel_HAv1 contigs were included in hybrid scaffolds. The hybrid scaffolds spanned 67 contigs, whereas the remaining 10 un-scaffolded contigs included most of those that were also incongruent with the genetic map. Six of the sixteen chromosomes were recovered as single scaffolds and each chromosome was represented by an average of 2.2 scaffolds.

We conducted additional scaffolding using the genetic map AmelMap3 [38] and Hi-C data, followed by gap filling and polishing in order to produce version Amel_HAv3. In this final version, each chromosome is represented by a single scaffold, comprised of an average of 4.2 contigs. Chromosomes 4 (13.4 Mbp) and 15 (9.5 Mbp) are recovered as single contigs, including the distal telomeres (see below). For comparison, in Amel_4.5, the chromosomes are comprised of 340 anchored scaffolds. Contigs are named after linkage group and order on the genetic map, i.e. Group1_2 for the second contig on linkage group 1. A full list of scaffolds, contigs and their length and placements is provided in **Supp. Table S1**. A visual overview of the 16 chromosomes is presented in **Fig 2**.

**Fig 2.**
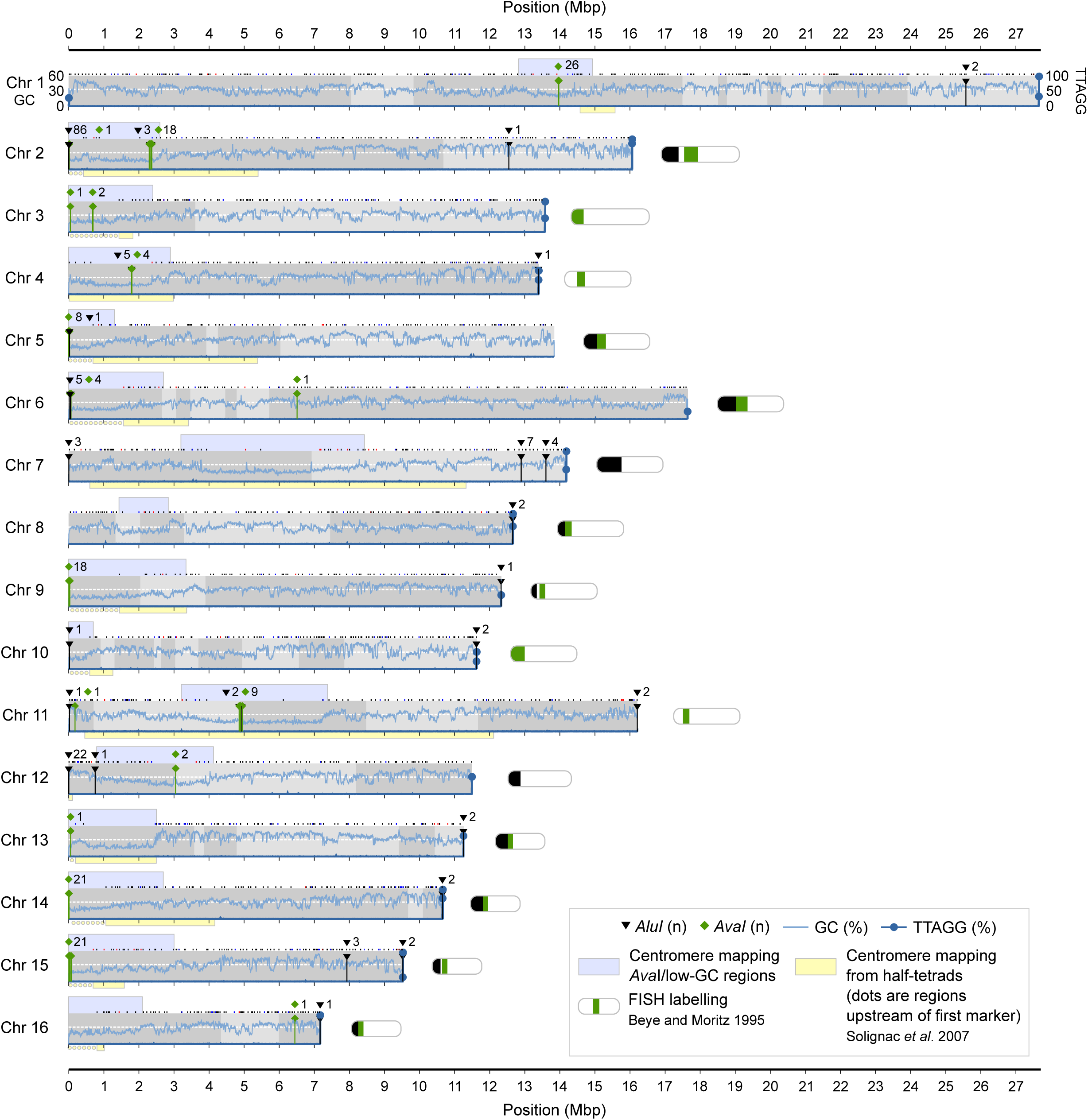
Assembly overview. An overview of the 16 linkage groups or chromosomes of Amel_HAv3 after anchoring and orienting the contigs according to the genetic map [38]. Grey shades indicate the intervals of each contig. Dots above each chromosome indicate the locations of genetic map markers (black=markers that are congruent with the assembly; red=markers that are incongruent, i.e. interleaved or reversed; blue=ambiguous markers, i.e. overlapping or widely separated primer sites). Genome-wide GC-content is indicated with a white dashed line and local %GC is mapped across all chromosomes (10 kbp non-overlapping windows; light-blue curve on y1-axis). The density of telomeric TTAGG/CCTAA repeats is shown (10 kbp non-overlapping windows; dark-blue curve on y2-axis; filled circles shown for values >10%). Extended low-GC regions indicating putative centromere regions are shown above chromosomes (bounded by adjacent 100 kbp windows < genome-wide %GC; light-blue), whereas experimental centromere mappings from [31] are indicated below chromosomes (boxes bounded by genetic map markers; extended upstream to the tip of the chromosome as dots when the area started at the first genetic map marker; light-yellow). The locations of centromeric *AvaI* (green) and telomeric *AluI* (black) clusters, respectively, are marked along chromosomes. Miniature chromosome models are redrawn from [30] and indicate experimental detection of *AvaI* and *AluI* arrays.

### Congruence of assembly with the genetic map

The order of genetic map markers in the linkage map AmelMap3 [38] was compared to their order on the Amel_HAv3 chromosome-length scaffolds. Out of a set of 4,016 paired primer sequences for 2,008 microsatellite markers (**Supp. Table S2**), we found that 301 primers for 268 markers did not map to the assembly (7.5% of primers; 13.3% of markers), including both primers for 33 markers. Thus 1,975 marker loci (98.4%) could be positioned along the chromosomes (avg. 123 markers per chromosome). Out of these, 1,885 (95.4%) are congruent and collinear between Amel_HAv3 and the genetic map and the scaffolds are nearly fully consistent with the order of contigs suggested by the genetic map (**Supp. Table S3)**. We find a small fraction (0.9%) of the markers to be ambiguous. The primer pairs were originally designed to amplify polymorphic microsatellites and are expected to map close together on the chromosomes and not overlap with other pairs. The BLAST targets were >1 kbp apart for only 10 primer pairs (0.5%) and for 8 pairs (0.4%) they were overlapping.

However, we also detected minor unresolved incongruences inside or between adjacent contigs. A total of 72 markers (3.6%) have inconsistent placements in Amel_HAv3. These include cases where a small number of adjacent markers were locally arranged in the opposite physical order along contigs, compared to the expected order in the genetic map or where markers from different adjacent contigs were mixed at their borders, producing interleaved or nested contigs with respect to their order in the genetic map. Removing markers at zero genetic distances to their adjacent markers (n=241) reduced this rate of inconsistency to 2.5%, suggesting that the original order of some of these markers in the genetic map is itself ambiguous. Interleaved/nested contigs were observed within 5 chromosomes: the 0.4Mbp contig Group6_2 appears to be partly discontinuous and nested within Group6_1 on chromosome 6; contig Group7_2 overlaps with the end of Group7_1 on chromosome 7; a single-marker from Group10_6 is associated with Group10_5 on chromosome 10; Group12_1 and Group12_2 are interleaved across a 0.1-0.2 Mbp region on chromosome 12; and a ~0.3Mbp segment of Group13_5 is found within Group13_6. These inconsistencies and marker primers that could not be placed on the new assembly may indicate unresolved assembly errors or other sequence differences around these microsatellite loci (e.g. missing or divergent target sequence between this assembly and that used to produce the markers). Alternatively, they may reflect natural structural variation between the sample used for this assembly and those used to produce the genetic map.

### Comparisons of anchored and unplaced contigs in Amel_4.5 and Amel_HAv3

The final hybrid assembly (Amel_HAv3) has 219.4 Mbp of contig sequence could be anchored to the 16 chromosomes, compared to 199.7 Mbp in the assembly Amel_4.5 (**Table 2**). The extra 19.7 Mbp distributed across the Amel_HAv3 chromosomes represents an increase of about 10%. In Amel_4.5, 87.2% of sequence is anchored to chromosomes, which are represented by an average of 20.6 scaffolds, whereas in Amel_HAv3, 98.0% of sequence is anchored to chromosomes, which are all represented by single scaffolds. After removal of unpolished/low coverage fragments, there are only 4.45 Mbp of unplaced contigs in Amel_HAv3 compared to 29.4 Mbp in Amel_4.5 and a substantial amount of sequence has effectively been transferred from previously unplaced scaffolds (see alignment analyses below). N50 of contigs anchored to linkage groups is 6.93 Mbp in Amel_HAv3 compared to 53 kbp in Amel_4.5 (131-fold improvement).

**Table 2.** Sequence content of hybrid assembly

| contig location | Amel4.5 |  |  | Hybrid assembly (Amel_HAv3) |  |  |
| --- | --- | --- | --- | --- | --- | --- |
|  | anchored | unplaced | all | anchored | unplaced | all |
| Size (Mbp) | 199.72 | 29.38 | 229.12 | 219.39 | 4.45 | 223.84 |
| Contigs (n) | 7,769 | 8,732 | 16,501 | 67 | 160 | 227 |
| Longest contig (Mbp) | 0.333 | 0.072 | 0.333 | 13.400 | 0.486 | 13.400 |
| Contig N50 (Mbp) | 0.053 | 0.006 | 0.046 | 6.930 | 0.037 | 5.381 |
| Contig L50 (n) | 1094 | 1152 | 1,390 | 12 | 24 | 13 |
| Scaffolds (n) | 340 | - | 340 | 16 | - | 16 |
| GC (%) | 33.98 | 23.94 | 32.70 | 32.72 | 23.45 | 32.53 |
| Repeats (%) | 6.80 | 10.51 | 7.27 | 7.50 | 21.55 | 7.78 |
| Mappability (avg. score) | 0.967 | 0.843 | 0.896 | 0.985 | 0.639 | 0.978 |
| Metazoa BUSCO genes (n,%) | 881 (90.1%) | 60 (6.2%) | 941 (96.2%) | 951 (97.2%) | 8 (0.8%) | 959 (98.1%) |
| Hymenoptera BUSCO genes (n,%) | 4088 (92.6%) | 222 (5.0%) | 4310 (97.6%) | 4322 (97.9%) | 15 (0.4%) | 4337 (98.3%) |

In Amel_4.5, 16.7 Mbp (7.3%) of the sequence is marked as repetitive and unplaced contigs have higher levels of repeat sequence than chromosome-anchored contigs (10.5% vs 6.8%; **Table 2**). In Amel_HAv3, the overall amount and proportion of repeats has increased to 17.4 Mbp and 7.9% (1.1-fold). In comparison to the overall addition of sequence to chromosomes (+10%; 219.4 in Amel_HAv3 vs. 199.7 Mbp in Amel_4.5), we find that repeat sequence has been added at twice this proportion (+21%; 16.5 Mbp vs. 13.6 Mbp), indicating that we have incorporated sequence with higher levels of repeats than the genomic background into chromosomes.

Several features distinguish contigs that we were unable to incorporate into the genetic map or scaffolds (**Table 2**). These contigs are lower in GC content, have a larger proportion of repetitive sequence and have lower mappability. These features are also present in Amel_4.5, but are more pronounced in Amel_HAv3. For instance, repeat content is 2.9-fold higher among the unplaced vs. anchored Amel_HAv3 contigs compared to 1.54-fold higher in Amel_4.5 (**Table 2**). These repeat sequences remain difficult to place even with current long-read technologies.

### BUSCO gene content

We compared the respective completeness of the Amel_4.5 and Amel_HAv3 assemblies by counting the number of universal single-copy orthologues detected in either assembly with BUSCO [39]. Overall, Amel_HAv3 has a slightly larger number of BUSCO genes compared to Amel_4.5 (98.3% compared to 97.6%) (**Table 2**). However, in Amel_4.5, 5-6% of these BUSCOs are detected among unplaced contigs, whereas only 0.4-0.8% of these occur in unplaced contigs in Amel_HAv3 (**Supp. Table S4**). The hybrid assembly therefore represents a significant improvement in terms of the proportion of conserved genes located in genome scaffolds.

### The mitochondrial genome

We recovered a complete mitochondrial genome (16,463 bp; %GC=15) and could detect and label all features along the sequence (13 coding genes; 22 tRNAs; 2 rRNAs) using a combination of BLAST [40] and MITOS [41]. All coding genes and rRNAs, and most tRNAs (n=15), were accurately detected using BLAST (<6 bp missing from canonical models). All tRNAs were detected near-full length using MITOS (<3 bp missing from canonical models). All features were found to be in full synteny with previous assemblies [34,42]. The Amel_HAv3 mitochondrial sequence is 120 bp longer than in these assemblies (16,463 bp vs. 16,343 bp). After aligning the sequences, we found that most of the length difference is explained by three major intergenic indels: i) a 16 bp deletion between COX3 and tRNA-Gly; ii) a 190 bp hyper-repetitive insertion in the AT-rich region (%AT=96.9) next to the small ribosomal subunit; and iii) a 39 bp deletion in the same region. The remaining 15 bp are due to small scattered 1-3 bp indels. The 190bp insertion was likely not possible to assemble before with Sanger or short-reads. The mitochondrial genome and structural variants are presented in **Supp. Fig 2** and feature coordinates are provided in **Supp. Table S5**.

### Repeat content

The honeybee genome has relatively few repeats compared to other insects (8%; Table 2). In both this and the previous assemblies (Amel_HAv3 and Amel_4.5), we find 12.8 Mbp of simple repeats/low complexity regions with RepeatMasker, representing 5.6% of the overall sequence and about 75% of all repeat-masked output (**Supp. Table S6**). The remaining share (5 Mbp) consists of longer interspersed DNA transposons, long/short interspersed nuclear elements (LINE/SINEs), long terminal repeats (LTRs), RNA sequences and other minor repeat classes. In agreement with previous analyses of transposable elements in honeybees [34], we find that DNA transposons are the major repeat class (3.1 Mbp; 66% of all interspersed repeats; 1.4% of the assembly; Fig 3A; **Supp. Table S6**), and that *mariner* transposons are the most common element within this class (1.74 Mbp; 56% of DNA transposons). Many repeats occur at approximately the same frequency in both assemblies under our analytical conditions (**Fig 3B-C**), although some repeat classes occupy larger proportions of the genome. For instance, DNA transposons are only 1.02 times more frequent (n=108/Mbp in Amel_HAv3 vs n=106/Mbp) but occupy 1.25 times more space (1.38% vs 1.11%) in Amel_HAv3 compared to Amel_4.5. Likewise, rRNA sequences occupy over two times as much sequence but occur at nearly the same frequency (**Supp. Table S6**). This discrepancy suggests that many repeat motifs are individually longer in Amel_HAv3 than in Amel_4.5.

**Fig 3.**
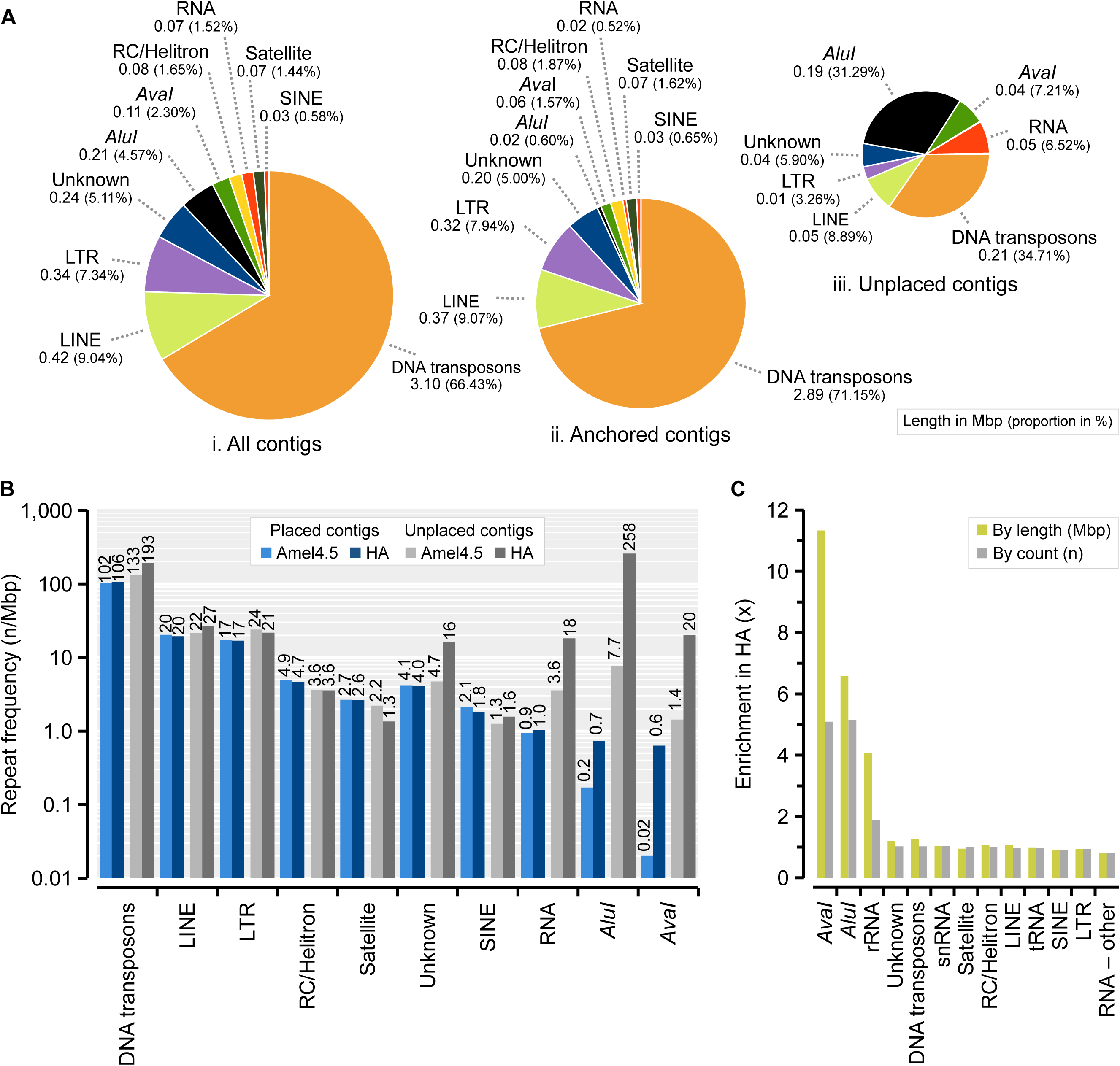
Interspersed and tandem repeats detected with RepeatMasker. A) The proportion of different repeat classes across the Amel_HAv3 in: i) all contigs; ii) anchored contigs; and iii) unplaced contigs. The total length and proportion of each repeat is given below each class. B) Comparison of repeat frequencies in anchored sequence and unplaced sequence between Amel_4.5 and Amel_HAv3. C) Overall enrichment of repeats in Amel_HAv3 compared to Amel_4.5.

The most striking difference in repeat annotation in Amel_HAv3 is the addition of a large number of *AvaI* (547 bp; n=229) and *AluI* (176 bp; n=1,315) repeats (**Fig 3B**; **Supp. Table S7**). These repeats have previously been estimated to represent 1-2% the honeybee genome using Southern blotting and FISH, and to be clustered close to centromeres (*AvaI*) and the short-arm telomeres (*AluI*) [30,43]. We detect 6.5 times more *AluI* repeat sequence in Amel_HAv3 than in Amel_4.5 (0.21 Mbp vs. 0.033 Mbp; n=1,315 vs. n=261) and 11 times more *AvaI* sequence (0.11 Mbp vs. 0.010 Mbp; n=229 vs. n=46; Fig 3C), although we are unable to fully assemble and map the complete sets (*AluI* abundance represents 0.14% of the assembly). Because many of the repeats occur in unplaced contigs (89% of *AluI* and 41% of *AvaI* repeats, respectively). The enrichment is lower by fragment count rather than overall sequence length (5.2-fold for *AluI* and 5.1-fold for *AvaI*). This is likely explained by higher repeat fragmentation in Amel_4.5, inflating repeat counts: only 30% (78 of 261) of *AluI* repeat matches are >160 bp in Amel_4.5, compared to 78% (1,022 of 1,315) in Amel_HAv3 (**Supp. Fig 3A**). Likewise, average divergence from the canonical *AluI* repeat is 15% in Amel_4.5 but only 3.9% in Amel_HAv3 (**Supp. Fig 3B**). For the *AvaI* repeats, only 2 % (1 of 46) are >500 bp in Amel_4.5 vs. 73% (167 of 229) in Amel_HAv3, and divergence is 21% vs. 6.6% (**Supp. Fig 3C-D**).

In Amel_HAv3, we find that *AluI* and *AvaI* repeats tend to cluster into extended tandem arrays (see **Fig 2** for their distribution on anchored contigs), often without any extra bases inserted between copies. The longest such anchored gap-free *AluI* array occurs on contig Group2_1 at the start of chromosome 2 and spans 80 adjacent full-length copies reiterated across 14.0 kbp (89 *AluI* repeats occur in the region; **Fig 4A**). Half of all *AluI* repeats occur in tandem arrays of at least 21 copies (~3.7 kbp). Two of these are at the short-arm ends of scaffolds (chrs. 2 and 12; **Fig 2**), while the rest are on unplaced contigs, indicating that these extensive *AluI* repeats are associated with the short-arm telomeres, as suggested previously [30,43]. In comparison, the longest contiguous *AluI* region in Amel_4.5 spans only 9 repeats (~1.4 kbp) on an unplaced contig. Likewise, the longest *AvaI* region in Amel_HAv3 occurs on chromosome 1 (contig Group1_3) and spans 26 copies across 14.1kbp (**Fig 4B**), and 50% of *AvaI* repeats occur in tandem arrays of at least 9 repeats (~4.4 kbp). No such gap-free *AvaI* array is detected in Amel_4.5. These improvements in Amel_HAv3 underscore the major advantage that long reads have over previous short-read technologies in resolving and representing highly repeated sequences.

**Fig 4.**
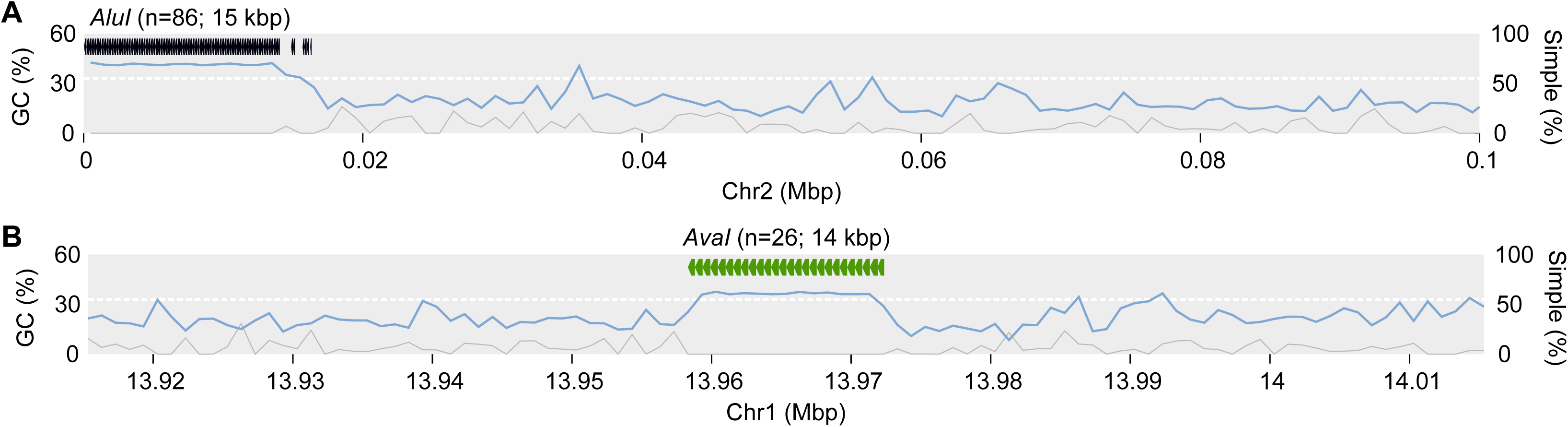
The Longest tandem arrays of *AluI* and *AvaI* repeats. A) Location of the longest *AluI* cluster. Genome-wide GC-content is indicated with a white dashed line and local %GC is shown across 1kbp non-overlapping windows (light-blue curve on y1-axis). Grey curve indicates the proportion of simple repeats (1kbp non-overlapping windows; y2-axis). B) Location of the longest *AvaI* cluster. Other statistics as in A.

### Alignment to previous builds

To further characterize differences between the genome assemblies, we aligned Amel_4.5 sequences against Amel_HAv3 using Satsuma [44]. Overall, alignments were produced against 94.3% of Amel_HAv3 (211.3/223.8 Mbp; see **Supp. Fig 3** for full alignment maps between assemblies). Chromosomal sequence was more frequently aligned (95.4% of 219.4 Mbp in Amel_HAv3) than unplaced contigs (44.4% of 4.45 Mbp in Amel_HAv3), which is consistent with these relatively repetitive contigs containing sequence that is not well represented in Amel_4.5. For sequences that had been associated with chromosomes in both assemblies, we found that 191.6 Mbp of alignments originated from the same chromosome in either assembly (99.4% of chromosome-to-chromosome alignments; 86% of all Amel_HAv3), while only a small fraction (1.23 Mbp; 0.56% of Amel_HAv3) originated from different chromosomes (**Fig 5A**), suggesting largely consistent mapping of data. About 16.4 Mbp of sequence that had previously been unplaced in Amel_4.5 now aligned against Amel_HAv3 chromosomes, corresponding to 7.5% of the total Amel_HAv3 assembly (**Fig 5A**). For comparison, we found that the opposite pattern was very uncommon: only 0.148 Mbp of alignments was mapped to chromosomes in Amel_4.5 but is unplaced in Amel_HAv3 (0.07% of Amel_HAv3). About 10.3 Mbp (4.7% of Amel_HAv3) was anchored to chromosomes but had no matching sequence in Amel_4.5 (conversely, 6.6Mbp of chromosomal sequence in Amel_4.5 is not matched in Amel_HAv3). Alignments were produced for 1.98 Mbp of contigs that are unplaced in both assemblies (0.9% of Amel_HAv3), whereas 2.32 Mbp unplaced Amel_HAv3 contigs did not align against Amel_4.5 (1.1% of Amel_HAv3; **Fig 5A**).

**Fig 5.**
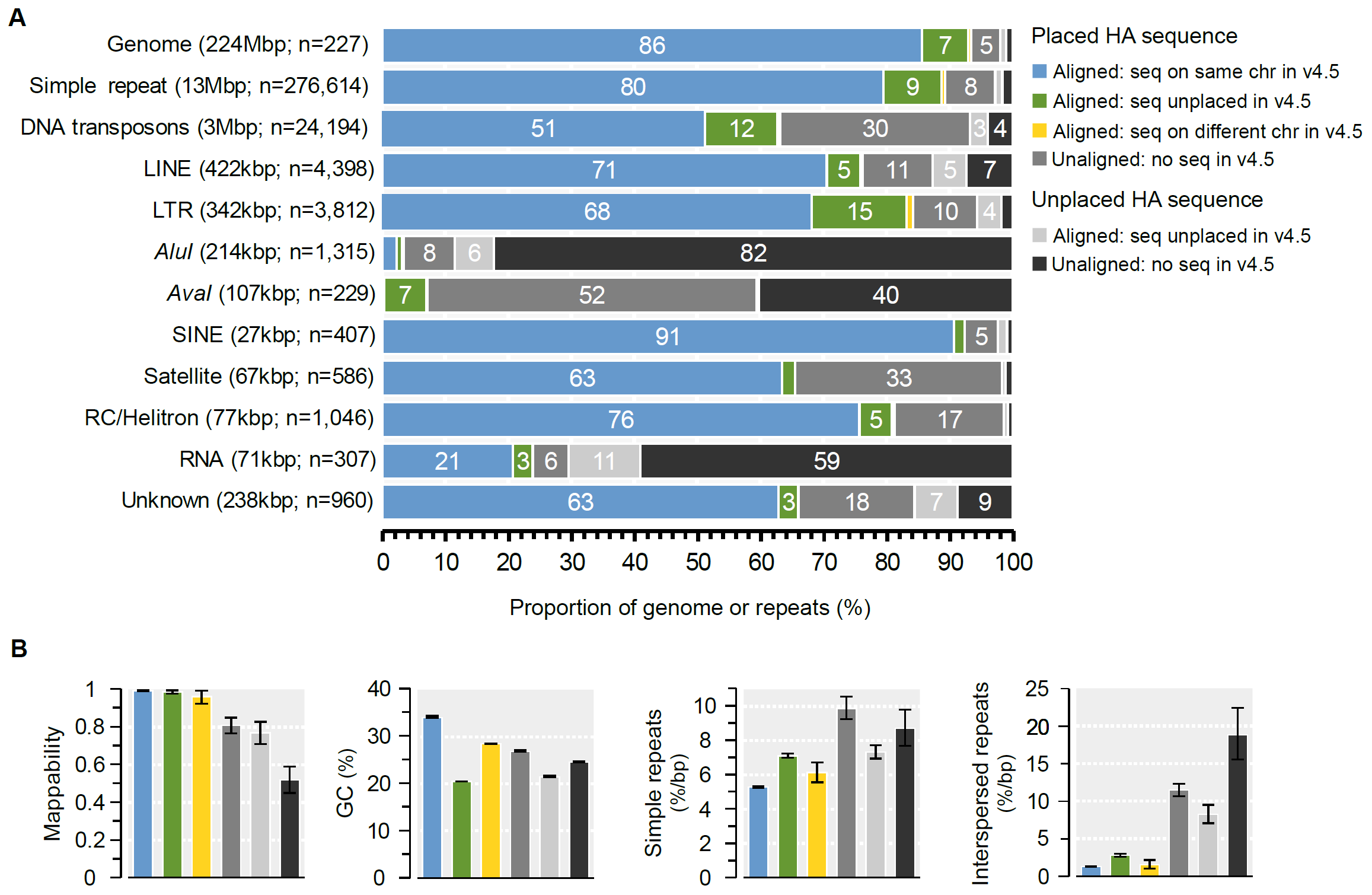
Properties of sequences classified from whole-genome alignments between Amel_HAv3 and Amel_4.5 using Satsuma. A) The proportions of the Amel_HAv3 assembly with or without matching sequence in Amel_4.5 is displayed at the top. The first four categories (left-to-right) refer to anchored sequence: blue=alignments between sequences that occur on the same chromosome in both assemblies; green=alignments between sequences that are anchored to chromosomes in Amel_HAv3 but were unplaced in Amel_4.5; yellow=alignments between sequences that have switched chromosomes; grey=unaligned Amel_HAv3 sequence without detected matches in Amel_4.5. The two last categories refer to unplaced sequence: light-grey=alignments between sequences that were not anchored to chromosomes in either assembly; dark-grey=unanchored and unaligned Amel_HAv3 sequence. The amount and proportion of simple repeats and the different classes of interspersed repeats according to the alignment regions in A is show below. B) The average mappability, %GC and density of simple and interspersed repeats/low complexity sequence according to the regions in A (95% confidence intervals generated from 2,000 bootstrap replicates of 1 kbp non-overlapping windows).

Aligned sequences that are anchored to the same chromosomes in either assembly have the highest average mappability scores (0.994) and GC content (34.1%; **Fig 5B**), characteristic for high-complexity/low-repeat sequence that is most amenable to assembly via last-generation technologies. Sequence that has been incorporated into chromosomes only in Amel_HAv3, but is unplaced in Amel_4.5 or unmatched/unaligned with Amel_4.5 sequence, has significantly lower GC-content (20.5% and 26.9%, respectively), and in the latter case also lower mappability (0.81). Both aligned and unaligned sequences that we are unable to place on chromosomes have reduced mappabilities and GC-content compared to genome genomic background (**Figure 5B**). Sequence that has switched chromosomes between assemblies has intermediate values for these statistics.

We find that newly anchored sequences or sequences that are still unplaced are significantly enriched for both simple and interspersed repeats (**Fig 5B**; see **Supp. Fig 5** for the density of individual repeat classes). Chromosomal regions built from sequences that were unplaced in Amel_4.5 (7%) or unaligned to Amel_4.5 sequence (5%), represents 12% of the genome but contain 17% of simple repeats (1.43-fold enrichment), 42% of DNA transposons (3.5-fold), 25% of LTRs (2.12-fold), 35% of satellites (2.93-fold) and 59% of *AvaI* repeats (4.94-fold; **Fig 5A**). Regional occurrence and enrichments for repeat-classes and their sub-classes can be found in **Supp. Table S8**.

### Distal telomeres

The telomeric repeat motif TTAGG is expected to occur as tandem arrays at the tip of the distal long-arm telomeres of all honeybee chromosomes. Distal telomeres were previously characterized from relatively short and fragmented sequences spanning only a few hundred base pairs of TTAGG repeats at the tips of five long-arm chromosomes in assembly Amel_4.0 [33], but manual scaffolding connected them to all but one long-arm chromosome tip [45]. We scanned for TTAGGs across 10 kbp windows in Amel_HAv3 and detected large clusters (on average 1,177 repeats; ~5.7 kbp) at the very ends of the long arms of 14 chromosomes (all except chromosomes 5 and 11; **Fig 2**; **Supp. Table S1**). While TTAGG/CCTAAs are rare across the genome (about 8 motifs per 10 kbp or ~0.4% of the genomic background; **Fig 2**), the outermost 1-2 windows of these chromosomes contain on average 1,043 motifs per 10 kbp (52% of the sequence; 130-fold enrichment; **Fig 2**). The longest telomeric repeat region was assembled for chromosomes 3 and 8, containing 2,142 and 1,994 copies of the motif, respectively. For the metacentric chromosome 1, we detected TTAGG repeats at both ends of the chromosome (the reverse complement motif CCTAA at the start of the chromosome), which is consistent with the hypothesis that this large chromosome has formed from fusion of two acrocentric chromosomes and harbors two distal telomeres [33,45].

We extracted and aligned the sequences of all distal telomeres with TTAGG arrays using MAFFT (n=15, including both telomeres on chromosome 1), including ~4 kbp of the upstream subtelomeric region, and scanned the sequences for shared properties. Taking the sequence at chromosome 8 as reference, we find that the first 2kbp downstream of the start of the telomere is enriched for TCAGG, CTGGG and TTGGG variants (**Fig 6A,C**). These polymorphisms are gradually replaced by the canonical TTAGG repeat moving towards the distal ends of the telomeres, where the average pairwise divergence between telomeres accordingly is much reduced: from 12% at <2 kbp away from telomere start to 2.4% at 2-4 kbp away (**Fig 6D**). We recover a relatively conserved 3 kbp subtelomeric region upstream of the junction (avg. pairwise divergence 14%; **Fig 6D**). The subtelomeres contain two larger shared motifs just upstream of the junction telomere junction (**Fig 6**): i) a ~350bp (213-520bp) fragment is located 100bp upstream of the junction and has moderate similarities towards a 4.5kbp LINE/CR1 retrotransposon originally characterized in *Helobdella robusta* (CR1-18_HRo; avg. ~74% identity; ii) a highly conserved and GC-rich 400 bp sequence (avg. div. 5.5%) is located further upstream but does not have significant similarities with any sequences in RepeatMasker or NCBI GenBank. Chromosomes 5 and 11 do not contain arrays of TTAGGs in Amel_HAv3, but terminate with subtelomere sequences that include the conserved motif (identified with BLAST). Three unplaced contigs contain a large number of motifs: the 18 kbp GroupUN_199 has 1,177 TTAGGs, the 16 kbp GroupUN_7 has 909 CCTAAs and GroupUN_198 has 82 CCTAAs.(**Supp. Table S1**). No other 10 kbp window contains >30 such motifs among the unplaced contigs. It is possible that these three contigs belong to the truncated chromosomes. Both GroupUN_7 and GroupUN_198 associate with chromosome 11 in the Hi-C dataset. GroupUN_198 also contains a >2.6 kbp subtelomeric subsequence (labeled with BLAST). Mate-pair reads with TTAGGs have previously been linked to the tip of chromosome 11 [45].

**Fig 6.**
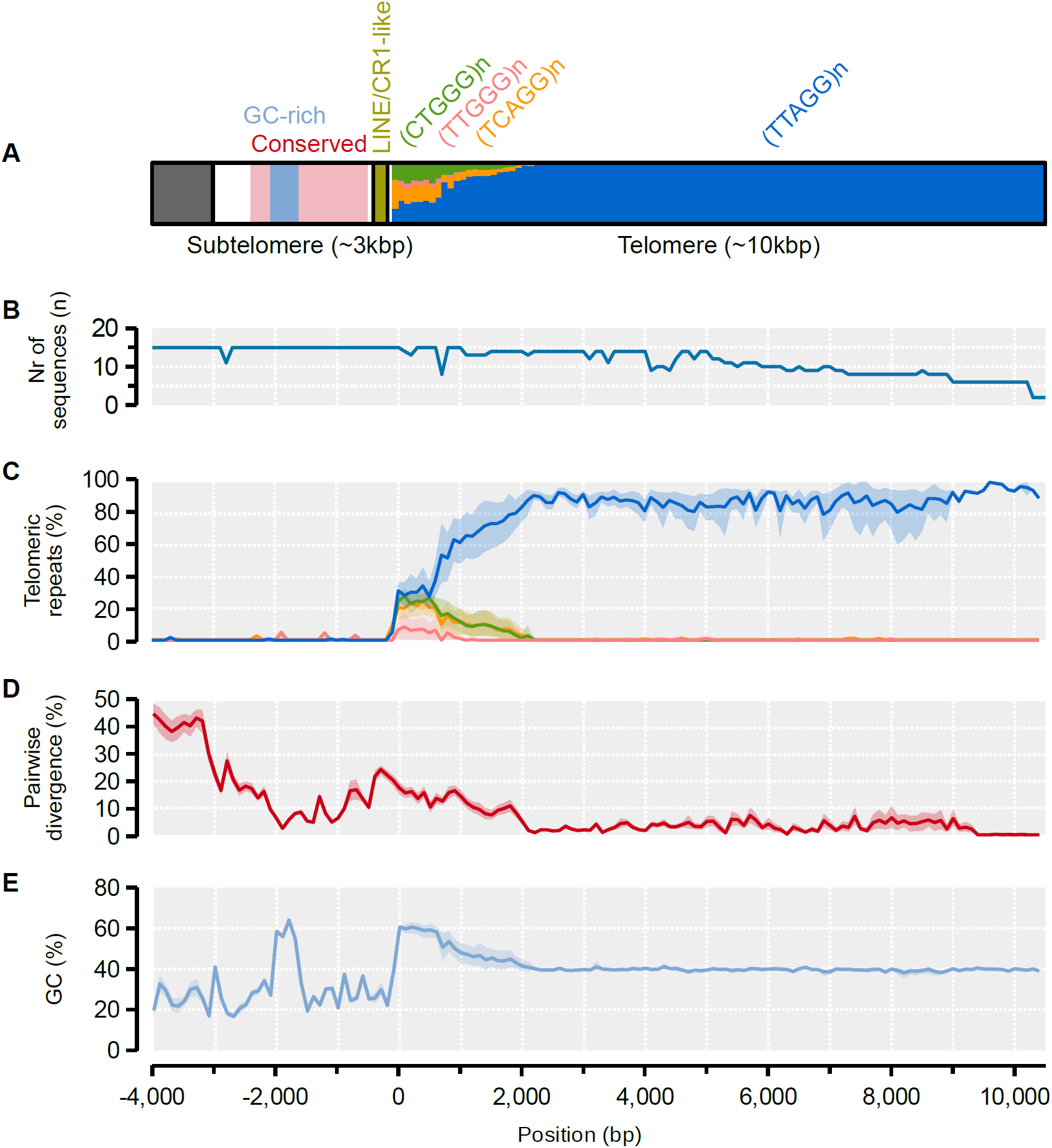
Model and properties of distal telomeres. A) A model of the subtelomeric and telomeric regions as inferred from alignment and sequence analysis of the distal ends of 14 chromosomes (two telomere sequences from chromosome 1). All statistics are computed across 100-bp windows using the distal telomere on chromosome 8 as backbone. A 3-kbp subtelomeric region is indicated with a white box, together with conserved and GC-rich sub-regions within it. A shared repeat element is indicated at the subtelomere-telomere junction. A >10-kbp telomeric region is indicated in the last box and the proportions of the canonical TTAGG repeat and variants are indicated for every 100-bp window. B) Number of subtelomere/telomere sequences extending across the alignment; C) The average density of TTAGGs and variants along the region. 95% confidence intervals for each window was computed from 2,000 bootstrap replicates. D) The average pairwise sequence divergence between chromosomes. Confidence intervals computed as in C. E) Average GC-content along the region. Confidence intervals computed as in C.

For comparison, we scanned Amel_4.5 for TTAGGs and subtelomeric sequence to locate telomeres in this assembly. In Amel_4.5, we find subtelomeres on short contigs (average length of 27kbp) located at the tips of the outermost scaffolds of 13 chromosomes. We detected TTAGG clusters with on average 34 motifs per telomere near five of these, whereas the rest had repeat densities that were indistinguishable from background levels. The distal telomere sequences in Amel_HAv3 are therefore 35 times longer (1,177/34) than those in Amel_4.5.

### Centromeres and proximal telomeres

*AvaI* and *AluI* repeats have previously been suggested to indicate the positions of centromeres and proximal short-arm telomeres, respectively, in the honeybee genome [30,43]. Although many *AvaI* and *AluI* repeats remain unmapped (see above), we find that the mapped repeats cluster toward the tips of the short-arms of most acrocentric (2-6, 9, 14-15) and the center of metacentric chromosome 1 and possibly submetacentric chromosome 11 (**Fig 2; Supp. Table S7**). These locations are largely similar to previous FISH labeling of these sequences for all but two chromosomes (10 and 16; **Fig 2**). We find that short-arm telomeric *AluI* repeats often co-occur with centromeric *AvaI* repeats (e.g. chromosomes 2, 4, 5 and 6). This is consistent with fluorescent labeling that also suggests that the proximal telomeres blend with centromeres in many acrocentric chromosomes [30]. However, the assembly often terminates at or near these clusters, sometimes before reaching into the proximal telomere (e.g. chromosomes 14 and 15; **Fig 2**).

The distribution of the putatively centromeric *AvaI* repeats in Amel_HAv3 overlaps or co-occurs with experimental mapping of centromeres from patterns of recombination and heterozygosity in half-tetrads of the clonal Cape honeybee *A. m. capensis* (e.g. chromosomes 2, 4, 11; **Fig 2**) [31]. The high contiguity in Amel_HAv3 now facilitates further characterization of the putative centromeric regions. All mapped *AvaI* clusters with more than two repeats (n=11; **Fig 2**) are embedded in megabase-scale regions with reduced GC content compared to the rest of the genome (22.7% vs. 34.6%; average length 2.3 Mbp; delineated by 100kbp windows with GC<32.7%; **Fig 2**). Sequences up to 1-2 Mbp away from the *AvaI* clusters have significantly reduced %GC and increased density of simple repeats and DNA transposons, compared to the genomic background (>2 Mbp away; p<0.05; 2,000 bootstrap replicates of data intervals; **Fig 7**). Patterns of centromeric enrichment were unclear for the rarer repeat classes. Similar low-GC blocks were detected in chromosomes 13 and 16, although only a single or no *AvaI* repeat, respectively, was mapped to these regions. The low-GC centromere-associated regions together span 42 Mbp of the genome and are among those that appear to have been particularly poorly assembled before: these regions constitute 19.3% of the genome but contain 38% of all sequence that is unmatched against Amel_4.5 (3.9 Mbp; 2.0-fold enrichment) and 95% of all sequence that was unplaced in Amel_4.5 (15.6 Mbp; 4.9-fold enrichment) (**Supp. Fig 4A**). These regions have more than doubled in size compared to Amel_4.5 (23.1 Mbp more than 19.2 Mbp; 2.2-fold increase).

**Fig 7.**
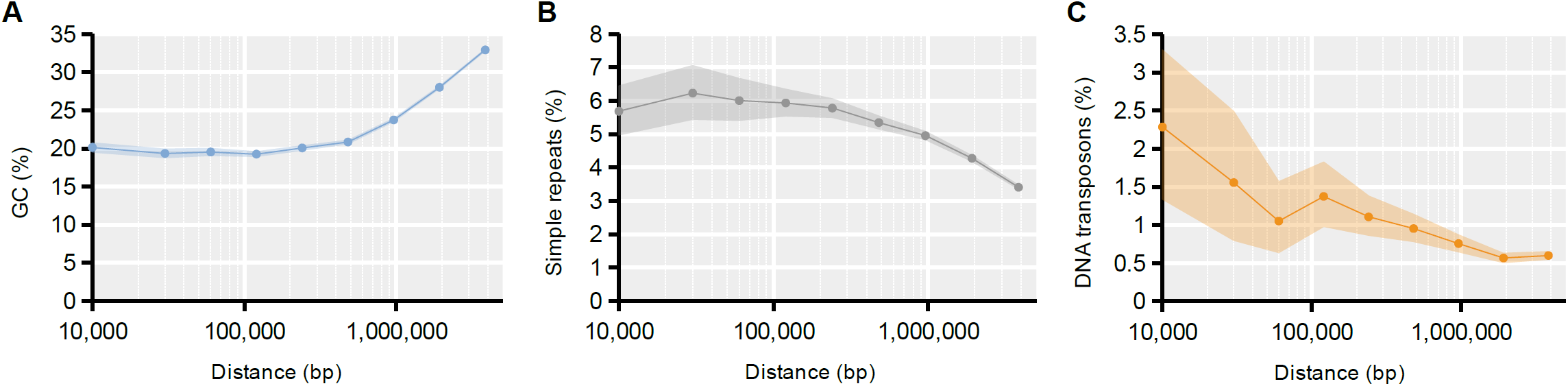
Features around centromeric *AvaI* repeats. A) Average GC-content was computed from 1kbp windows located within intervals at different distances from *AvaI* clusters with at least 3 repeats (0-20kbp; 20-40kbp; 40-80kbp; 80-160kbp; 160-320kbp; 320-640kbp; 640-1,280kbp; 1,280-2,560kbp; 2,560-5,120kbp). 95% confidence intervals were computed from 2,000 bootstrap replicates of each interval. B) As in A but tracing the density of simple repeats/low complexity sequence. C) As in A, but tracing the density of DNA transposons, the dominant interspersed repeat class in the honeybee genome.

We next used the genetic distances previously inferred between the genetic map markers to compare recombination rates inside and outside of these regions. Across the genome, we estimate the average recombination rate to be 21.6 cM/Mbp (n=1,735 congruent marker pairs), close to what has been estimated before in honeybee [38,46,47]. Compared to these background levels, recombination rates are significantly reduced across both sets of centromere mappings: to 14.6cM/Mbp (0.68-fold; n=249) in the half-tetrad experiment from [31] and to 7.9cM/Mbp (0.38-fold; n=99) from our assessment of *AvaI* and GC-content (**Fig 8**). In contrast to the FISH results, we also detect several small *AluI* clusters close to long-arm telomeres (**Fig 2**). However, compared to the repeats at the proximal telomeres, these hits are fewer (32 vs. 130), shorter (106 bp vs. 162 bp) and more divergent (16% vs. 4.5%) on average (**Supp. Fig 3A-B**), which could indicate excess spurious hits or degenerate elements.

**Fig 8.**
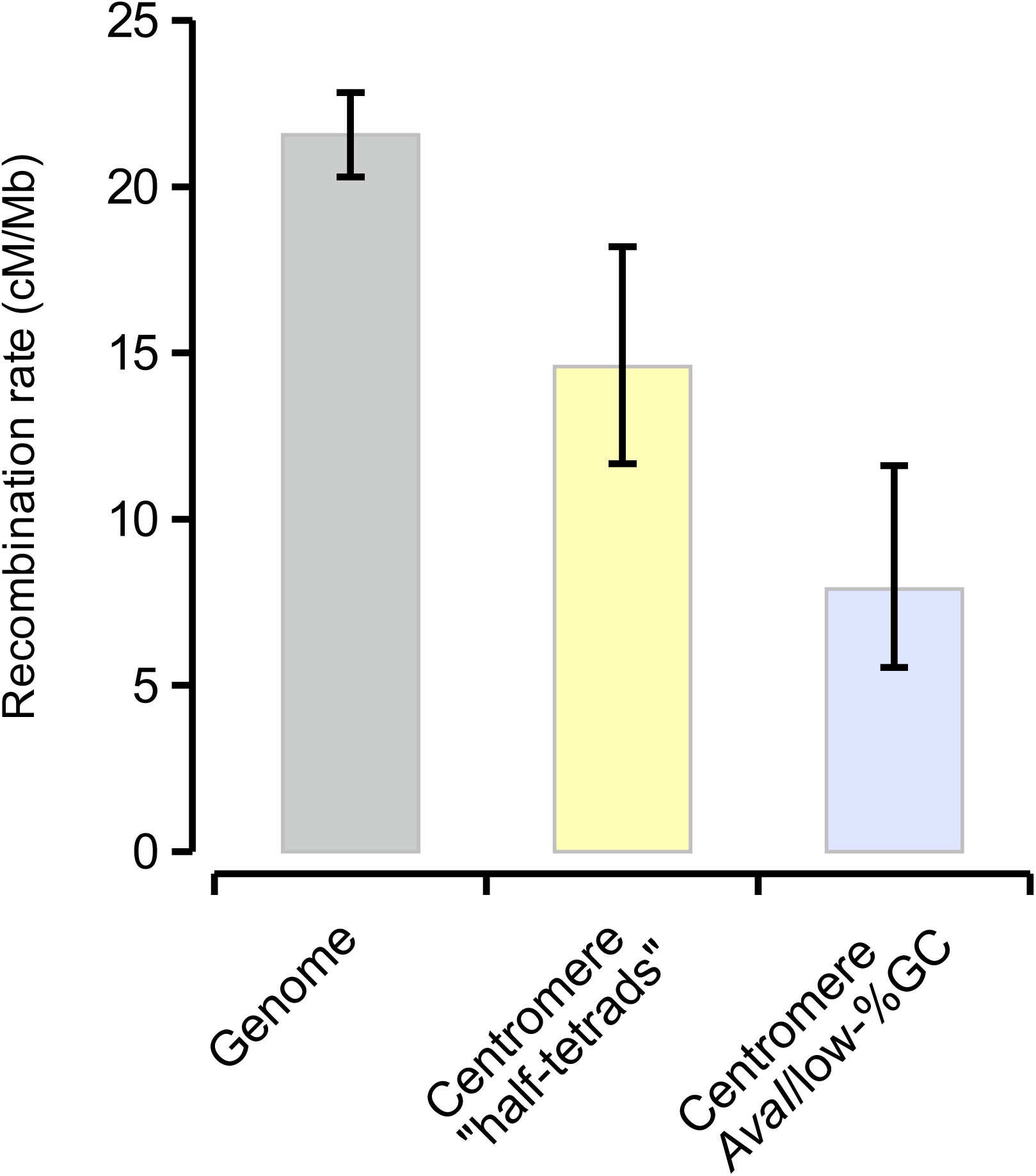
Recombination rates in different genomic regions. Recombination rates were computed from the genetic and physical distances between genetic map markers scattered across the whole genome or located within putative centromere regions. 95% confidence intervals were computed from bootstrapping marker-to-marker pairs (2,000 replicates).

We do not find TTAGGs associated with proximal telomeres, suggesting they are either not present at the short-arms of the honeybee chromosomes or only occur in unmappable sequence. To address this, we manually inspected mate-pairs sequenced from decade-old fosmid libraries that were prepared for the original assembly [33]. Fosmid reads containing *AluI* repeats were found to likely have *AluI* mate pairs, indicating very long strings of *AluI*s that supersede the length of the arrays in the hybrid assembly. Interestingly, out of 19 mate pairs containing the TTAGG motifs and linked back to telomere regions, only 7 anchored to distal telomeres, while 12 contained *AluI* repeats. This suggests they belong to proximal telomeres, although no individual read contained directly observable junctions between *AluI* and TTAGG motifs. This independent evidence nevertheless suggests the presence of TTAGG repeats beyond the currently mappable regions of *AluI* repeats on the short-arm telomeres. Moreover, the CCTAA enriched unplaced contig GroupUN_7 (see above) also contains 28 *AluI* repeats, and could potentially be a proximal or mis-joined contig. Because our assembly of these regions between the centromeres and the short-arm telomeres remains incomplete, most of the unplaced contigs are inferred to belong in these regions.

## Discussion

Here we have produced a hybrid assembly (Amel_HAv3) for the Western honeybee using PacBio long-reads merged with 10x Chromium linked-reads to generate extremely long contigs. These contigs were scaffolded using a BioNano optical map, a Hi-C chromatin conformation map, and the genetic linkage map AmelMap3 [38]. This pipeline enabled us to produce a highly contiguous genome (contig N50=5.4 Mbp; scaffold N50=11.3 Mbp). In Amel_HAv3, there are on average 4.2 contigs per chromosome and two of sixteen chromosomes (4 and 15) are recovered near end-to-end as single contigs. All chromosomes are reconstructed as single scaffolds. The assembly represents a 120-fold improvement in contig-level contiguity and 14-fold scaffold-level contiguity, compared to the previous assembly Amel_4.5 (Fig 1; Table 1).

The assembly was constructed using an incremental approach, where each step resulted in linking or scaffolding existing contigs and thus extending contiguity. This is currently the only approach possible for combining multiple technologies and generating a hybrid assembly. It is beneficial to construct long contigs prior to scaffolding to accurately align them to optical maps or chromatin conformation data. However, assembly errors that incorrectly join sequence are possible at each step and increases in contiguity may come at the expense of freezing errors into the assembly. This entails a tradeoff between completeness, contiguity and accuracy. Ideally, an approach that integrates all technologies simultaneously to weigh and minimize conflicts between different approaches to construct the optimal assembly is needed although no such methods currently exist [48].

We identified several instances of conflicts between the assembly and scaffolding technologies used, which emphasizes the value of using multiple sources of data. In particular, the availability of a genetic linkage map was crucial to evaluate such conflicts. However, there are still 2-3% markers in the genetic map that do not align colinearly with our assembly. These regions require further evaluation, but one likely explanation is that our assembly and the genetic map are based on different strains of honey bee. Consistent with other reports [23] we find that our Hi-C data were highly accurate at assigning contigs to linkage group but resulted in orientation errors and placement errors, revealed by comparison with the genetic map and BioNano scaffolds. We therefore only used these data to assign and confirm assignment to linkage group. A particular advantage of the honeybee for genome assembly is their haplodiploid mode of sex determination which results in the availability of haploid (male) drones, which eliminates the difficulties posed by heterozygous sites.

We have incorporated ~10% more sequence into chromosomes compared to Amel_4.5 (more than the full length of a typical honeybee chromosome). The newly anchored sequence has low GC-content and high repeat content. Much of this sequence can be traced to previously unplaced fragments in Amel_4.5, and as a consequence, most unplaced single-copy orthologues have now been transferred to chromosomes. Many repeat classes occur at approximately the same frequencies between the hybrid assembly and Amel_4.5, but for several classes (including DNA transposons, rRNA sequences and centromeric/telomeric repeats) we detect appreciably longer matches against the canonical database motifs (**Fig 3; Supp. Table S6**). This suggests higher accuracy in assembling these repetitive elements with the new sequencing technologies deployed here, compared to the Sanger and short-read sequences used for the Amel_4.5 assembly [34]. The hybrid assembly contains substantially more repetitive sequence comprising both centromeres and telomeres than the previous one, which unifies the assembled chromosome sequences with the karyotype as observed under the microscope [30,33] (**Fig 2**). However, the longest tandem repeat arrays associated with these features are about 14-15 kbp (**Fig 4**), less than 10% of their experimentally inferred size (see below) and are likely limited by the upper read-lengths of our PacBio libraries.

Most of the new sequence incorporated into this genome assembly compared with the previous one is anchored as Mbp-scale blocks of low-GC heterochromatin around the centromeres of most chromosomes. These regions make up about 19% of the genome and are enriched for repetitive sequence and DNA transposons (**Fig 7**). In agreement to what has been shown in many other taxa [20,49], we find that these centromeric regions have reduced rates of meiotic recombination (**Fig 8**). Honeybee centromeres have been shown to contain extended arrays of the 547 bp *AvaI* repeat that appears to make up about 1% of the genome (~300 repeats across 150 kbp per centromere) using Southern blotting and FISH [30]. It was not possible to demonstrate an association between *AvaI* and centromeres in previous assemblies due to the relative absence of the *AvaI* repeat and poor contiguity of these regions [33,34]. The scaffolds in Amel_HAv3 are highly congruent with the genetic linkage map AmelMap3 [38] and the *AvaI* repeats typically coincide with the expected location of centromeres based on linkage maps [31].

Honeybee telomeres have two different structures. Short-arm telomeres (which are close, or proximal, to the centromeres) consist of tandem arrays of the 176bp *AluI* element that make up as much as 2% of the genome (~2,000 repeats or 350kbp per telomere), as estimated with restriction enzymes and fluorescent probes [30,43]. Telomeres on the long arms of chromosomes (distal to centromeres) have shared subtelomeric blocks that are followed by extended iterations of the TTAGG repeat and were originally characterized along with the first published honeybee assembly [33,45]. The TTAGG repeat is likely ancestral for insect telomeres [50–52] and has been estimated to range between 2-48kbp in size among chromosomes using Southern hybridization [53]. The difference between proximal and distal telomeres has been hypothesized to support chromosome polarity and pairing during cell division [45].

Our hybrid assembly contains repeat arrays associated with both proximal and distal telomeres. Although TTAGG repeats may be present beyond the *AluI* arrays on the short-arm telomeres, we are unable to conclusively map any TTAGGs to this end of the chromosomes and only anchor them to the distal telomeres on the long arms. Here they stretch up to 10kbp beyond the subtelomere, within the expected size range for honeybee telomeres [53]. Close to the junction between the subtelomeres and the telomeres, we recover a large number of variant motifs (**Fig 6**). About 90% of the TCAGG and CTGGG variants co-occur in the higher order repeat TCAGGCTGGG, which has also been detected in previous assemblies [54]. The origin of this diversity is unclear, but their localization towards the inner telomere suggests they are older more degenerate sequences compared to the more homogenous sequence of the outer telomere.

A major utility of a highly contiguous genome assembly is that it can be used as a basis to reveal structural variants such as inversions, duplications and translocations that are obscured in more fragmented genome assemblies [7]. Structural genomic variation is an important source of phenotypic variation, and it is crucial to survey this form of variation in order to identify genetic variants associated with gene regulation, phenotypic traits or environmental adaptations [55]. Breakpoints of structural variants are commonly associated with repetitive elements that often reside in gaps in more fragmented assemblies. A striking example of adaptation likely governed by structural variation is observed in high-altitude populations of *A. mellifera* in East Africa, where highland and lowland populations are highly divergent in two distinct chromosomal regions [56]. In species of *Drosophila* fruit flies, a large number of cosmopolitan chromosomal inversions have been identified that govern adaptation to environmental clines [57]. Notable examples of inversions that govern environmental adaptation have also been found in stickleback fish and *Heliconius* butterflies [58,59]. It is therefore likely that much important phenotypic variation maps to structural variation in honeybees and that this contiguous genome assembly will be an important resource for uncovering it.

### Conclusion

We have produced a highly complete and contiguous genome assembly of *A. mellifera* by combining data from four long-read sequencing and mapping technologies. The strength of this hybrid approach lies in combining technologies that work at different scales. PacBio data consist of long (>10 kb) reads but it is problematic to incorporate extended repetitive regions into contigs assembled from these data. We therefore used linked-read 10x Chromium data to bridge gaps between contigs and fill them with additional sequence data. Long contigs produced by this approach could then be scaffolded effectively by BioNano optical mapping and Hi-C chromatin conformation mapping to result in chromosome-length scaffolds. The assembly is particularly improved in repetitive regions, including telomeres and centromeres. This new genome sequence assembly will facilitate research into the functioning of these regions and into the causes and consequences of structural genomic variation.

## Methods

### Library preparation and data production

We produced data using Pacific Biosciences SMRT sequencing (PacBio), 10x Chromium linked-read sequencing (10x), BioNano Genomics Irys optical mapping (BioNano) and a Hi-C chromatin interaction map (Phase Genomics). DNA extracted from a single drone pupa from the DH4 line was used for the first three of these methods (a different drone for each method). These individuals were brothers of the individuals from the DH4 line used for previous honeybee genome assembly builds [33,34]. The sample used for Hi-C was an individual from an unrelated managed colony with a similar genetic background (mixed European) as the DH4 line (mixed European) collected from the USDA-ARS Bee Research Laboratory research apiary.

To prepare DNA for the PacBio and 10x sequencing, we first lysed cells from 20-120 mg of insect tissue. This was done by grinding in liquid nitrogen followed by incubation at 55°C in cell lysis solution (25 ml 1 M Tris-HCl, pH 8.0; 50 ml 0.5 M EDTA, pH 8.0; 0.5 ml 5 M NaCl; 12.5 ml 10% SDS; 162 ml molecular grade water) and proteinase K. The solution was then treated with RNase A. Proteins were then precipitated using Protein Precipitation Solution (Qiagen) and centrifugation at 4°C. DNA was precipitated from the resulting supernatant by adding isopropanol and ethanol and centrifugation at 4°C.

We generated a 10 kb PacBio library that was size-selected with 7.5 kb cut-off following the standard SMRT bell construction protocol according to manufacturers recommended protocols. The library was sequenced on 29 SMRT cells of the RSII instrument using the P6-C4 chemistry, which generated 10.2 Gb of filtered data. N50 subread length was 8.8 kb. A 10x GEM library was constructed from high-molecular-weight DNA according to manufacturers recommended protocols. The resulting library was quantitated by qPCR and sequenced on one lane of a HiSeq 2500 using a HiSeq Rapid SBS sequencing kit version 2 to produce 150 bp paired-end sequences. This resulted in 127,440,953 read pairs (38Gb of raw data).

High-molecular-weight DNA was extracted *in situ* in agarose plugs from a single drone pupa following BioNano Genomics guidelines. Plugs were cast and processed according to the IrysPrep Reagent Kit protocol with the following specifications and modifications; a 7-day proteinase K treatment in lysis buffer adjusted to pH 9.0 with 2µl BME per ml buffer. The BspQI NLRS reaction was processed according to protocol, stained overnight and immediately loaded on 2 flow cells for separation on the BioNano Irys system. In total 1,214,651 molecules were scanned with N50 of 210 kbp.

DNA for the Hi-C experiment was prepared at Phase Genomics. The sample was incubated at 27°C for 30 minutes with periodic mixing by inversion. Glycine was added (final concentration of 0.1g/10mL) to quench crosslinking. After an additional incubate at 27°C for 20min with periodic inversion, the sample was pelleted by centrifugation, the supernatant was removed and the sample was kept at −20°C prior to processing. This procedure results in extraction of native cross-linked chromosomes from bee cells, disruption using endonucleases and linking of adjacent strands via biotinylated junctions. The samples were then sequenced on an Illumina HiSeq instrument.

### Assembly pipeline

In order to determine the best way to utilize the data, we first generated assemblies using the PacBio and 10x Chromium data independently (see below). As the PacBio assembly had far superior contiguity, we designed a pipeline to begin with this assembly and then use the 10x linked-reads to connect and combine contigs. These contigs were then scaffolded using the BioNano optical map data, with additional checks for consistency with Hi-C data and a genetic map [38]. Gap filling and polishing steps were also included to maximize contiguity and accuracy. Full details of the pipeline are presented below and summarized in **Supp. Fig 1**.

We imported PacBio raw data into the SMRT Analysis software suite (v2.3.0) (Pacific Biosciences, CA) and generated subreads. All sequences shorter than 500 bp or with a quality (QV) <80 were filtered out. The resulting set of subreads was then used for *de novo* assembly with FALCON v0.5.0 [60] using pre-assembly length cutoff of 7 kbp. Since the genomic DNA originated from haploid drone we kept only primary contigs generated by FALCON and removed 14 contigs shorter than 2kbp before further analysis. The resulting set of contigs was polished twice using Quiver via SMRT Analysis Resequencing protocol [60]. The resulting PacBio assembly consisted of 429 contigs with N50 of 3.1Mbp and largest contig being 9.7Mbp.

To create the 10x Chromium assembly we used Supernova 1.1.4 on the 10x Chromium linked read data [61] with default parameters. The resulting assembly had 9734 scaffolds with N50 of 0.59Mbp and the longest scaffold was 3.2Mbp. This assembly was not used to create the final assembly and we instead used the 10x linked-read data to extend the PacBio assembly generated in the previous step. We ran the ARCS+LINKS Pipeline [35,36] to utilize the barcoding information contained in 10x linked reads. First, we mapped 10x reads to PacBio contigs using LongRanger 2.1.2 (10X Genomics, CA). ARCS v1.0.1 was then used to identify pairs of contigs with evidence that they are connected based on the observation of linked reads from the same molecule. Default parameters were used, except for modifying barcode read frequency range (-m 20-10000). The results of ARCS were processed with the LINKS v1.8.5 scaffolding algorithm to constructs scaffolds based on 10x read pairing information. We adjusted the –a parameter, which controls the ratio of barcode links between two most supported graph edges, to 0.9. The ARCS+LINKS pipeline produced 299 scaffolds with N50 of 8.8Mbp and longest scaffold of 13.3Mbp.

We compared the the PacBio+10x assembly to the genetic linkage map AmelMap3 [38] by determining the position of 2,008 microsatellite markers using BLAST [40,62]. This enabled us to assign 49 scaffolds to one of 16 linkage groups. The remaining sequences were designated as unplaced. Furthermore, we used genetic map information to order, orientate and to join adjacent scaffolds belonging to the same linkage group by introducing arbitrary gap of 2000 Ns. We then used PBJelly from PBSuite v15.8.24 [37] to perform a first round of gap filling using all PacBio reads. PBJelly closed 87 (67%) gaps within scaffolds due to joins made by ARCS+LINKS and 16 (48%) of the gaps that were introduced between adjacent scaffolds on the basis of proximity according to the genetic map. In order to minimize possibility of freezing scaffolding errors we then split scaffolds on remaining gaps. This stage of assembly resulted in assembly version Amel_HAv1 which had 330 contigs with N50 of 5.6Mbp and longest contig of 13.4Mbp.

The BioNano raw data were assembled using the BioNano Solve (v3.1.0) assembly pipeline (BioNano Genomics, CA) on a Xeon Phi server resulting in 171 genome maps with cumulative length of 285 Mbp and N50 of 2.2 Mbp. This data set was combined with Amel_HAv1 by running the BioNano Solve v3.1.0 hybrid scaffolding pipeline. The BioNano software identified 7 conflicts between optical maps and Amel_HAv1. All of the conflicts could be traced back to original FALCON assembly and were confirmed to be chimeric. Therefore, we chose to resolve these conflicts in favor of the BioNano optical maps. The resulting hybrid assembly had N50 of 11.3Mbp and a longest scaffold of 27.7Mbp length. This version of the assembly was designated Amel_HAv2.

Hi-C read pairs were aligned to the initial assembly using BWA-MEM [63] with the "-5" option. Unmapped and non-primary alignments were excluded using samtools [64] with the "-F 2316" filter. We next performed scaffolding with Hi-C data using the Proximo pipeline (Phase Genomics), which builds on the LACHESIS scaffolding package [14]. In total 149 out of 280 Amel_HAv2 scaffolds could be grouped into 16 clusters out which 14 clusters contained scaffolds that were previously assigned to linkage groups and each had cumulative size of predicted chromosome length. Two clusters contained short contigs without linkage group assignments. Two large Amel_HAv2 scaffolds were not a part of any Hi-C cluster, most likely due to fact that they already represented complete chromosomes. Overall, we observed completely accurate assignment of scaffolds to chromosomes based on comparison with genetic map. However, there were a number of errors in orientation and order of scaffolds within a Hi-C cluster. In total there were 22 such errors affecting 8 Hi-C clusters. Therefore, in the final chromosome scale scaffolds we ordered and orientated Amel_HAv2 scaffolds based on genetic map information. Scaffolds from the same linkage group and same Hi-C cluster were joined by introducing an arbitrary gap of 200 Ns in the orientation and order indicated by the genetic map.

We performed an additional round of gap filling using PBJelly to fill gaps generated by the previous scaffolding steps using all PacBio reads, which closed 12 (20%) additional gaps within the scaffolds. New sequences introduced during gap filling steps originate from non-error corrected subreads. To remove potential sequencing errors, the whole assembly was once more subject to two rounds of Quiver polishing. Subsequently 89 unplaced contigs that had fewer than 50% of their bases polished were removed from the final assembly. In addition, two unplaced contigs were identified as mitochondrial and removed. The final assembly consisted of 16 chromosomal scaffolds with a total of 51 gaps, 160 unplaced contigs and a mitochondrial sequence. This final data set was designated as hybrid assembly Amel_HAv3.

### Assembly characterization and analysis

After the scaffolding was completed, the congruence between the genetic map and the final assembly was reassessed. The primer sequences for the microsatellite markers in AmelMap3 were again fitted against the genome using BLAST. The physical positions and order of the markers along and between contigs was compared to their expected order in the linkage map.

The assembly was scored for base composition, mappability and repeat content. These metrics may correlate with sequences that are challenging to assemble. We compared the chromosome-anchored and unplaced sequences of the published reference assembly (Amel_4.5; [34]) to the new hybrid assembly (Amel_HAv3) for these properties. We computed average GC content across whole assemblies, arbitrary regions and along non-overlapping windows of different sizes (1 kbp; 10 kbp). We then used GEM v1.315b [65] to compute the mappability (or uniqueness) of short non-degenerate (0 bp mismatch) 50bp kmers across the assemblies. Every base is annotated for an average mappability score computed from overlapping kmers. We computed average mappability scores across windows as above.

We used BUSCO v2.0.1 [39] to compare the completeness of Amel_4.5 and Amel_HAv3 by assessing the number of expected and detected single-copy orthologs in either assembly as inferred from the OrthoDB v9.1 [66]. Two core sets of BUSCOs (near-universal single-copy orthologs) were used: Metazoa (n=978 BUSCOs) and Hymenoptera (n=4,415 BUSCOs).

We used RepeatMasker v4.0.7 [67] to annotate simple and interspersed repeat content. We deployed the RMBLAST-NCBI search engine to scan for animal repeats (-species metazoa) in the 20170127 release of the Repbase database [68]. The query and database was extended (-lib) to include the consensus motifs of two tandem repeats associated with centromeres (*AvaI*; 547bp; X89539) or proximal short-arm telomeres (*AluI*; 176bp; X57427), respectively, of honeybee chromosomes [30,43]. These elements were named after the bacterial restriction endonucleases originally used to detect them (AvaIR from *Anabaena variabilis*; AluIR from *Arthrobacter luteus*) but are unrelated to the similarly named Ava and Alu SINE class repeats of other taxa and not detected using the Repbase database. The canonical *AvaI* repeat consists of four highly similar sub-repeats, resulting in spurious overlapping annotations when *AvaI* repeats occur in tandem. We therefore parsed the pre-ProcessRepeats output (ori.out-file) separately to extract non-overlapping *AvaI* repeats. Simple repeats and low-complexity sequence as annotated by RepeatMasker were considered together.

In order to locate distal telomeres, we estimated the density of the short telomeric repeat motif TTAGG/CCTAA [45] across 10kbp windows in both Amel_HAv3 and Amel_4.5 using a custom Perl script. Distal telomeric and subtelomeric regions (<5kbp upstream of putative telomere start) were then extracted and aligned with the L-INS-i algorithm in MAFFT v7.310 [69]. The sequence in chromosome 8, which has among the longest telomeric repeat regions, was taken as a profile and columns with gaps in this sequence was removed from the alignment. The average pairwise sequence divergence, GC-content and density of TTAGG and variant repeats (TCAGG, CTGGG, TTGGG) was then estimated across 100bp windows along the alignment and 95% confidence intervals were computed by bootstrapping the sequences (n=2,000 replicates). We searched Amel_4.5 for a conserved subtelomere sequence that was shared between chromosomes in Amel_HAv3 (see Results) to help locating telomeres in this assembly using BLAST [40,62]. Lastly, we queried Amel_HAv3 RepeatMasker output for shared interspersed repeats among the subtelomeric regions.

Satsuma v2 [44] was used to align Amel_4.5 against Amel_HAv3 using default settings. The alignments were used to characterize the sequences found to be shared between the assemblies, or unique to either of them. The Amel_4.5 sequence is not oriented with respect to the genetic map and we did therefore not perform in-depth assessments of synteny or reorientations between assemblies.

The mitochondrial genome sequence was recovered in a single contig. It was circularized and subject to two rounds of polishing. We then used BLAST to detect the order and orientation of the canonical set of coding sequences, rRNAs and tRNAs along the chromosome (NCBI accession NC_001566) [34,42]. Because BLAST did not label all tRNAs in its default settings, we also used MITOS [41] together with MIFTI [70] to annotate the mitochondrial genome *ab initio*. The sequence was visualized in DNAPlotter [71]. In order to detect any major structural differences, we used MAFFT to align the whole mitochondrial sequence against the corresponding sequence in Amel_4.5.

## Declarations

*Ethics approval and consent to participate*

Not Applicable

*Consent for publication*

Not Applicable

*Availability of data and material* The *Apis mellifera* whole genome shotgun project has been deposited at NCBI under the accession QIUM00000000 and BioProject ID: PRJNA471592.

### Competing interests

The authors declare that they have no competing interests

### Funding

This research was supported by grants 2013-722 from the Swedish Research Council Formas and 2014-5096 Swedish Research Council Vetenskapsrådet to MTW. HMR was partly supported by a Romano Professorial Scholarship and funding from Greg Hunt. Driscoll’s Berries provided partial support for Hi-C sequencing via Phase Genomics. The funding bodies played no role in the design of the study and collection, analysis, and interpretation of data or in writing the manuscript.

### Authors’ contributions

The genome assembly and analyses were conducted by AW and IB. The manuscript was written by AW, IB and MTW with contributions from all other authors. The study was conceived by GER, HMR, and MTW. OVP, M-BM, AKC, JDE and ASM contributed data and were involved in project planning. The optical mapping experiment was performed by M-BM. All authors read and approved the final manuscript.

## Acknowledgements

We thank Alvaro Hernandez (Carver Biotechnology Center, University of Illinois) for assistance with the Chromium library preparation and 10x Genomics sequencing and Ivan Liachko and Shawn Sullivan (Phase Genomics) for assistance with the Hi-C data. The PacBio sequencing and BioNano optical mapping were performed at the Uppsala Genome Centre. Computation was performed at UPPMAX (Uppsala Multidisciplinary Center for Advanced Computational Science).

## Supplementary information

### Supplementary Figures

**Supp. Fig 1.** Assembly pipeline. Flowchart illustrating the assembly process. Data sources used as input are displayed in cyan, methods are displayed in yellow, and assembly versions are displayed in green. The final assembly, version 3, is designated Amel_HAv3

**Supp. Fig 2.** A map of the mitochondrial sequence in the hybrid assembly (Amel_HAv3). A) Summary statistics are presented in the center of the circularized sequence, followed by a 100bp sliding-window (20bp steps) bar-plot of GC-content relative to the mitochondrial average (15%). Major structural indels between Amel_HAv3 and Amel_4.5 mitochondrial sequences are indicated as black boxes. The order and orientation of the coding genes (pink), rRNAs (green), tRNAs (blue) are illustrated as arrows. The AT-rich region is indicated in deep purple. Coordinates are given in the outer circle. B) Alignments between Amel_HAv3 and Amel_4.5 illustrate base-level coordinates and composition of the structural variants highlighted in A.

**Supp. Fig 3.** Properties of *AluI* (176 bp) and *AvaI* (547 bp) RepeatMasker matches in the hybrid assembly (Amel_HAv3) and Amel_4.5. A) The length distribution of masked *AluI* repeats in either assembly. These are further subdivided according proximal or distal ends of chromosomes in Amel_HAv3. B) The distribution of sequence divergence from the canonical *AluI* motif. Classes and colors as in A. C) The length distribution of *AvaI* matches. D) The distribution of sequence divergence from the canonical *AvaI* motif. Classes and colors as in C.

**Supp. Fig 4.** Genome-wide Satsuma alignments between hybrid assembly (Amel_HAv3) and Amel_4.5. A) Alignments across every chromosome. Upper plot: genome-wide GC-content is indicated with a white dashed line and local %GC is mapped across all chromosomes (10kbp non-overlapping windows; light-blue curve on y1-axis). The density of telomeric TTAGG repeats is shown on the y2-axis (10kbp non-overlapping windows; dark-blue curve with circles). Average GEM mappability scores is show on y2-axis (10kbp non-overlapping windows; grey curve). Lower plot: Amel_4.5 scaffolds (upper grey arrows) aligned against Amel_HAv3 contigs (lower black arrows). Coordinates are Mbp-scale. Colors indicate aligned blocks (blue=alignments between sequences that occur on the same chromosome in both assemblies; green=alignments between sequences that are anchored to chromosomes inAmel_HAv3 but were unplaced in Amel_4.5; yellow=alignments between sequences that have switched chromosomes). White spaces are unaligned regions. The locations of centromeric *AvaI* (green) and telomeric *AluI* (black) clusters, respectively, are marked along chromosomes. B) As in A, but for unplaced fragments.

**Supp. Fig 5.** Density of repeat elements in different genomic regions in the hybrid assembly (Amel_HAv3). Density of interspersed and tandem repeats in different Amel_HAv3 regions, with or without matching sequence in Amel_4.5 (see Fig 5A for detailed definitions). 95% confidence intervals were generated from bootstrapping randomly extracted blocks of 1 kbp.

### Supplementary Tables

**Supp. Table S1.** Contigs and chromosomes in Amel_HAv3.

**Supp. Table S2.** Hybrid assembly (Amel_HAv3) contig-level coordinates of genetic map markers from AmelMap3 [38].

**Supp. Table S3.** Summary of the congruence and conflict observed between the hybrid assembly (Amel_HAv3) and the genetic map markers from AmelMap3 [38].

**Supp. Table S4.** BUSCOs detected in Amel_HAv3 and Amel_4.5.

**Supp. Table S5.** Features located in the mitochondrial sequence of Amel_HAv3.

**Supp. Table S6.** Repeat density in Amel_HAv3 and Amel_4.5.

**Supp. Table S7.** *AluI* and *AvaI* repeats in Amel_HAv3 and Amel_4.5.

**Supp. Table S8.** Repeat density across different alignment regions between Amel_HAv3 and Amel_4.5.

